# Acrylamide Fragment Inhibitors that Induce Unprecedented Conformational Distortions in Enterovirus 71 3C and SARS-CoV-2 Main Protease

**DOI:** 10.1101/2020.11.06.370916

**Authors:** Bo Qin, Gregory B. Craven, Pengjiao Hou, Xinran Lu, Emma S. Child, Rhodri M. L. Morgan, Alan Armstrong, David J. Mann, Sheng Cui

## Abstract

RNA viruses are critically dependent upon virally encoded proteases that cleave the viral polyproteins into functional mature proteins. Many of these proteases are structurally conserved with an essential catalytic cysteine and this offers the opportunity to irreversibly inhibit these enzymes with electrophilic small molecules. Here we describe the successful application of quantitative irreversible tethering (qIT) to identify acrylamide fragments that selectively target the active site cysteine of the 3C protease (3C^pro^) of Enterovirus 71, the causative agent of hand, foot and mouth disease in humans, altering the substrate binding region. Further, we effectively re-purpose these hits towards the main protease (M^pro^) of SARS-CoV-2 which shares the 3C-like fold as well as similar catalytic-triad. We demonstrate that the hit fragments covalently link to the catalytic cysteine of M^pro^ to inhibit its activity. In addition, we provide the first demonstration that targeting the active site cysteine of M^pro^ can also have profound allosteric effects, distorting secondary structures required for formation of the active dimeric unit of M^pro^. These new data provide novel mechanistic insights into the design of EV71 3C^pro^ and SARS-CoV-2 M^pro^ inhibitors and identify acrylamide-tagged pharmacophores for elaboration into more selective agents of therapeutic potential.

## INTRODUCTION

RNA viruses cause significant morbidity and mortality in human and animal hosts.^1,2^ For example, Enteroviruses (EV) include many important human pathogens with the best characterised being enterovirus 71 (EV71), rhinovirus (HRV), coxsackievirus B3 (CVB3), enterovirus D68 (EV-D68) and poliovirus (PV). EV71 is one cause of hand, foot and mouth disease (HFMD) in humans and is associated with severe neurological disease with considerable mortality.^3^ Vaccines against EV71 have been developed and approved ^4^ but outbreaks persist and there are no antiviral drugs available for treating EV71.^5,6^

Like many other RNA viruses, EV71 relies on proteases to cleave a polyprotein precursor into individual functional mature proteins. For enteroviruses, the majority of this proteolytic processing utilises the 3C protease (3C^pro^).^7^ Given the essential role of virally-encoded proteases in viral life-cycles, numerous protease inhibitors have been developed for potential clinical use.^8^ These include a large collection of picornaviral 3C^pro^ inhibitors, such as an HRV 3C^pro^ inhibitor Rupintrivir (AG7088)^9^ that failed to show patient benefit in phase II clinical trials.^10^ To date, no 3C^pro^ inhibitors have been approved for clinical use.

With the global rise of COVID-19, scientific attention has focussed on the causative agent, SARS-CoV-2.^11^ This virus expresses two precursor polyproteins (pp1a and pp1ab) that are cleaved by both main protease (M^pro^)^12^ and papain-like protease (PL^pro^).^13^ Interestingly, M^pro^ and 3C^pro^ share a similar fold, active site architecture and catalytic triads, both proteases being absolutely reliant on the catalytic cysteine for activity.^14^ Exploiting these structural similarities may enable identification of pharmacophores that target a wide variety of viral proteases.

Therapeutics with a covalent mechanism of action are becoming more widely accepted for a range of diseases.^15^ Targeted covalent therapeutics such as Ibrutinib^16^ and Osimertinib^17^ use a Michael acceptor to react with the thiolate of cysteine, giving irreversible target engagement.^18^ Identifying the starting points for such agents is often the bottleneck in development. Fragment-based approaches offer an efficient starting point and have already shown significant promise in targeting SARS-CoV-2 M^pro^.^19^ We have recently developed quantitative irreversible tethering (qIT), a high-throughput method for identifying selective covalent fragments that bind to a desired cysteine on a target protein.^20^ qIT enables hit prioritization and minimization of false positives and negatives through normalization of the rate of protein modification by compound intrinsic reactivity.^21^

Here we employ qIT to identify inhibitory fragments that covalently target the active site cysteine of EV71 3C^pro^. Cocrystals of 3C-fragment complexes demonstrated the occupancy of a novel, cryptic pocket in 3C^pro^. Furthermore, when repurposed towards M^pro^, the covalent fragments also preferentially targeted the active site cysteine, inhibiting the enzyme activity and, in one case, additionally disrupting the quaternary structure of M^pro^.

## RESULTS AND DISCUSSION

### Covalent fragment screening against EV71 3C^pro^ by quantitative Irreversible Tethering (qIT)

To target cysteine C147 on EV71 3C^pro^, we constructed a 1040-member covalent fragment library using a combination of in-house parallel synthesis and commercial vendors (Figure 1A and 1B). Fragment-like core scaffolds were functionalized with cysteine-reactive chemical groups, with the majority (>95%) being acrylamides. Acrylamide “warheads” are featured in several clinically approved covalent drugs and are favoured for their mild electrophilic reactivity, minimizing potential non-specific reactivity and associated toxicity.^22^ In line with the generally accepted FBDD guidelines, the library was designed to maximise scaffold diversity and to conform to the ‘rule of 3’:^23^ MW < 300, clogP ≤ 3, H-bond donors/acceptors ≤ 3 (Figure 1C).

**Figure 1.**
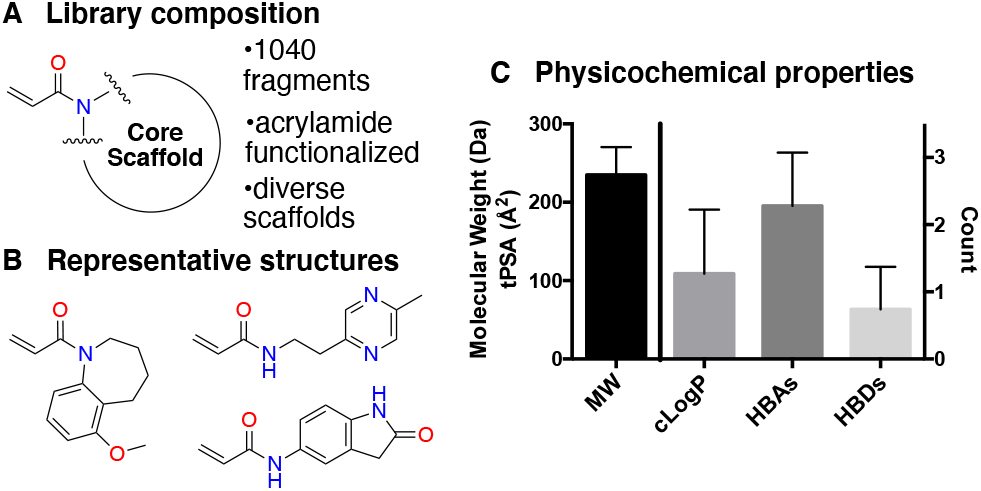
Cysteine-reactive covalent fragment library composition. (A) Covalent fragment library composition. (B) Representative structures of library members. (C) Physicochemical properties of the library: MW = molecular weight; HBA = hydrogen-bond acceptor; HBD = hydrogen-bond donor.

We applied our fluorescence-based covalent fragment screening platform (qIT) to identify fragments which covalently bind to C147 on EV71 3C^pro^ (Figure 2A). To determine the rate of reaction between a cysteine thiol and an acrylamide fragment, the cysteine quantification probe CPM is employed to measure the degree of cysteine modification at a series of timepoints. The CPM probe competes with the acrylamide for modification of the cysteine residue such that the fluorescence signal is inversely proportional to the extent of acrylamide-cysteine labelling, allowing the rate of reaction (*ν*) to be determined by exponential regression analysis in high-throughput. Our workflow uses GSH as a control cysteine-containing biomolecule and hit fragments are those that react significantly faster with 3C^pro^ than with GSH (Figure 2B). The selectivity of the fragment towards the 3C^pro^ is quantified by the rate enhancement factor (REF) which was used to identify and prioritise hit fragments (Figure 2C).

**Figure 2.**
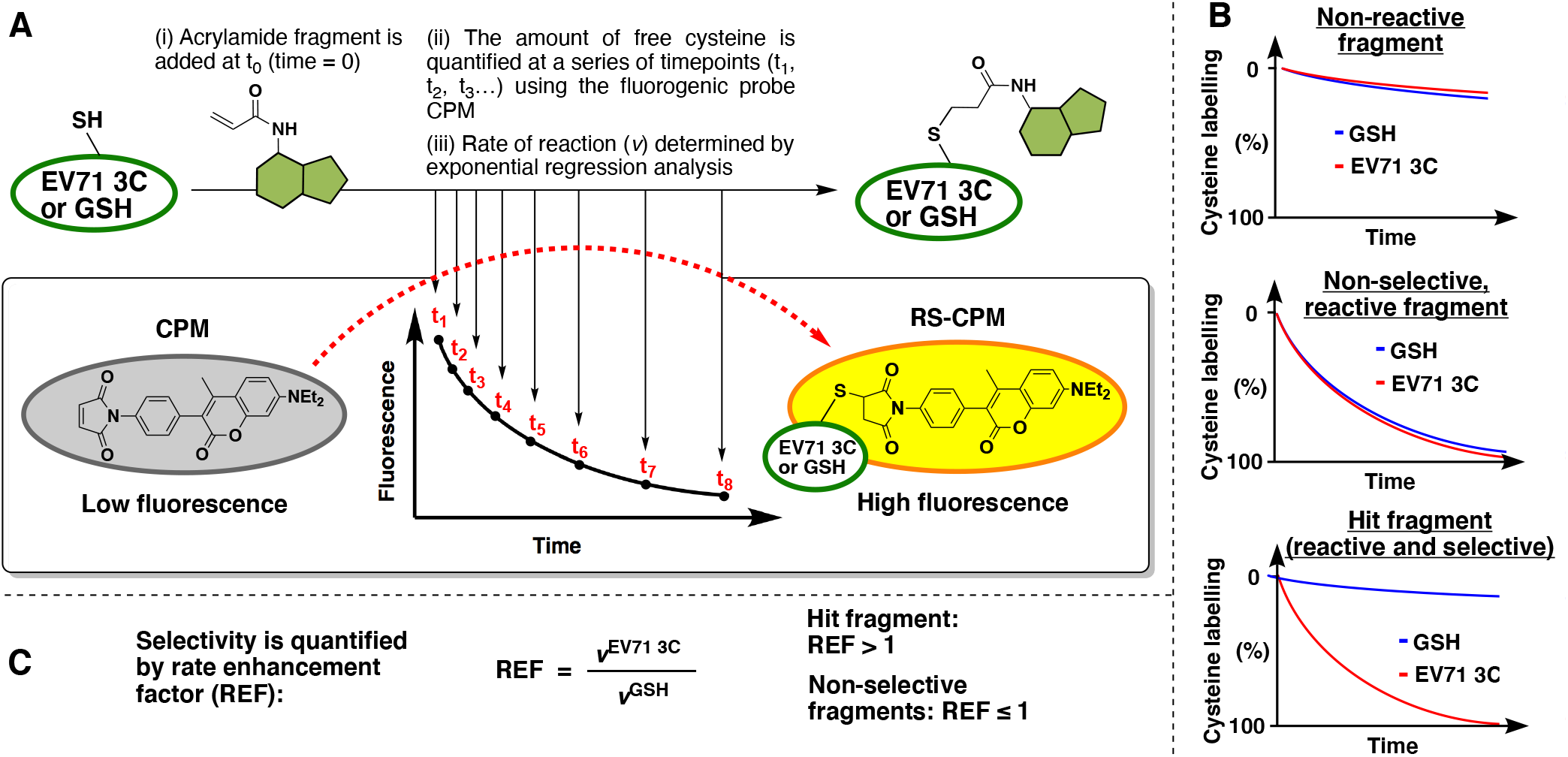
Quantitative Irreversible Tethering (qIT) screening platform. (A) Assay overview: The target thiol (5 μM), EV71 3C^pro^ or glutathione, is reacted with acrylamide fragments (0.5 mM) under pseudo-first order conditions. Reaction progress is followed by discrete measurements of free target thiol concentration using the fluorogenic probe CPM and the rate of reaction (v) are derived from exponential regression analysis. (B) Fluorescence intensity is converted into percentage cysteine modification by normalizing to DMSO control = 0%, no thiol = 100%. Fragments are characterized as (i) non-reactive, (ii) reactive but non-selective or (iii) reactive and selective by comparing the reactivity profiles between EV71 3C^pro^ and GSH. (C) Kinetic selectivity is quantified by the rate enhancement factor (REF) which is used to identify and prioritise hit compounds.

The 1040-member acrylamide fragment library was screened at 500 μM against EV71 3C^pro^ (5 μM) or GSH (5 μM) in parallel and the fluorescence intensity measured over 24 hours. The majority of the library (61%) displayed measurable reactivity with EV71 3C^pro^ over the 24-hour time course, with roughly half of those fragments showing selectivity over GSH (REF > 1) (Figure 3A). There were 13 fragments which had a REF greater than three standard deviations (1SD = 2.5) over the geometric mean (geomean REF = 1.0), for example acrylamide **1**, and these were taken forward for repeat qIT testing and mass spectrometry validation (Figure S1). Pleasingly, four of those fragments both had reproducible qIT profiles and clearly mono-modified EV71 3C^pro^ by intact protein mass spectrometry (Figures 3B, 3C and S2). The glutathione selectivity of the four validated hit fragments ranged from REF = 8.5–24.3 and the compounds shared some common chemical features: Acrylamides **1** and **2** both contain the same 1,3-thiazole core and methylene linker while acrylamides **3** and **4** have similar isoxazole motifs with more extended linkers. Encouragingly, all 4 hits are alkyl acrylamides which typically are associated with low levels of off-target reactivity and this is supported by their slow reactivity with glutathione.^24^

**Figure 3.**
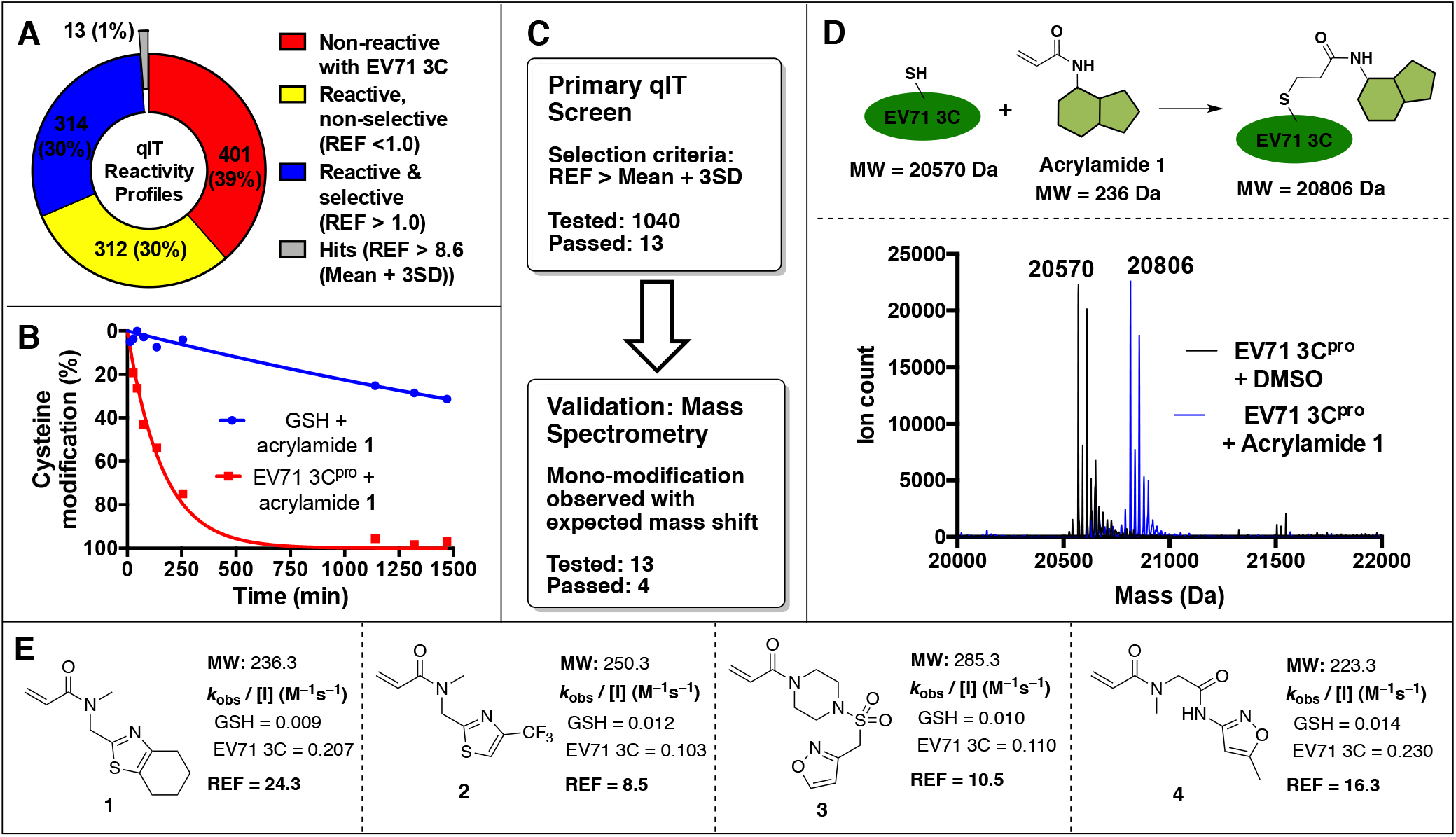
Screening cascade and hit validation. (A) Primary qIT screen summary: Pie chart shows number of fragments characterized as non-reactive, reactive but non-selective or reactive and selective. Hits have REF > 3 standard deviations over the geometric mean. (B) Illustrative qIT data for acrylamide **1** (0.5 mM) in reaction with EV71 3C^pro^ or glutathione (5 μM). (C) Summary of screening cascade. (D) Illustrative intact protein mass spectrometry data for modification of EV71 3C^pro^ (5 μM) reacting with acrylamide **1** (0.5 mM for 750 minutes). (E) Structures and reaction rates for the four validated hit fragments.

### The hit fragments covalently bind to residue C147 of EV71 3C^pro^ and accommodate a novel cryptic pocket

To reveal the binding site of the cysteine-reactive fragments, we labelled recombinant EV71 3C^pro^ with acrylamide fragments **1**–**4** and subjected the resulting complexes to crystallization trials. Unfortunately, the WT EV71 3C^pro^-fragment complexes did not yield crystals so we employed a 3C^pro^ mutant construct bearing the H133G to expedite the structural studies. The H133G mutant has WT-level protease activity, containing a WT-like catalytic triad, and harbors the H133G mutation at the hinge region of the β-ribbon, which improves the flexibility of the β-ribbon.^25^ Using the H133G mutant, we were able to determine crystal structures of 3C^pro^-**1** and 3C^pro^-**2** complexes; however, the structure determination of the other complexes remained unsuccessful.

Crystal structures of 3C^pro^-**1** and 3C^pro^-**2** complexes were solved to the resolution of 1.2–1.3 Å respectively, which provided atomic details of the 3C^pro^-fragment interactions. We found that both fragments **1** and **2** bind to the same pocket on 3C^pro^, with the acrylamide functionalities forming covalent bonds to C147 (bond length C(acrylamide)–S(C147)= 1.8 Å). Unexpectedly, neither of the fragments occupy the central substrate pocket (S1-S4) of 3C^pro^, as is observed for AG7088 and other characterized 3C^pro^ inhibitors.^9^ Instead, the fragments accommodate a cryptic pocket on the other side of the catalytic cysteine, denoted the S’ pocket. To obtain bias-free structural insight, we calculated the composite omit maps for fragments **1** and **2**, which clearly show that the fragments are buried inside the S’ pocket and defines the orientation of the thiazole rings with the sulfur atoms contacting the bottom of the pocket (Figure 4A & 4C). Comparing the binding of fragments **1** and **2**, we find that the trifluoromethyl and cyclohexane functionalities make less contact with protein and are associated with weaker electron density, hinting that the thiazole ring is the key pharmacophore.

**Figure 4.**
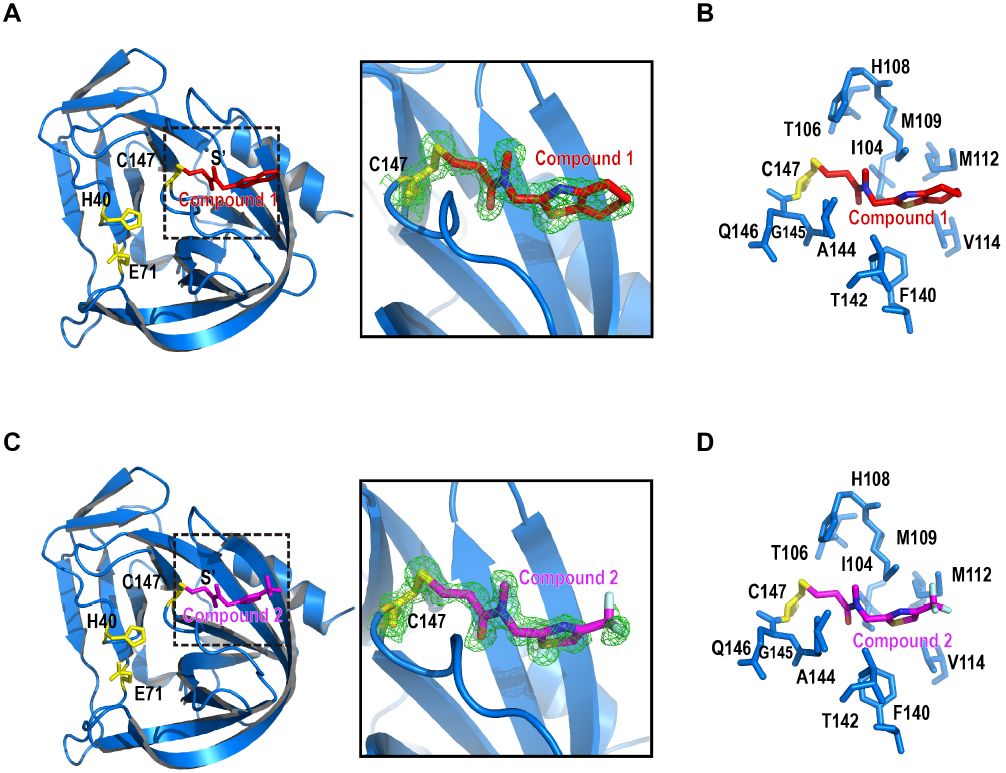
Crystal structures of EV71 3C^pro^-cysteine reactive fragment complexes. (A) Ribbon model of EV71 3C^pro^ (H133G) covalently linked to fragment **1**. Catalytic triad C147-H40-E71 (yellow) and fragment **1** (red) are shown with stick model. Right, magnified view of the dashed line box on the left. A composite omit map (contour level = 1.5) is superimposed with the stick model of fragment **1**. (B) Stick model of fragment **1** (red) binding the cryptic S’ pocket (blue) identified on the leaving group side of EV71 3C^pro^ active site. (C) Ribbon model of EV71 3C^pro^ (H133G) covalently linked to fragment **2**. Catalytic triad (yellow) and fragment **2** (magenta) are shown with stick model. Right, magnified view of the dashed line box on the left. A composite omit map (contour level =1.5) is superimposed with the stick model of fragment **2**. (D) Stick model of fragment **2** (magenta) binding the cryptic S’ pocket (blue) identified on the leaving group side of EV71 3C^pro^ active site.

**Figure 5.**
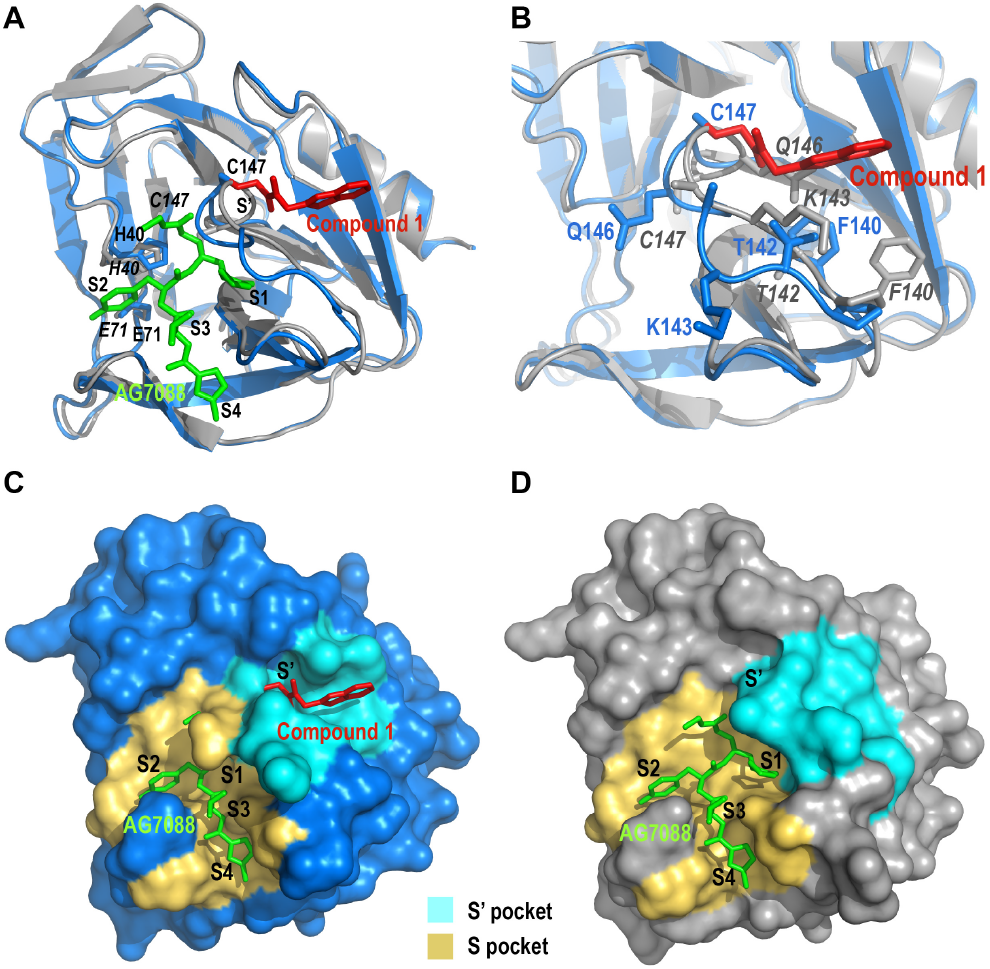
Fragment binding induces significant conformational rearrangement at the active site and substrate pockets. (A) The structure of EV71 3C^pro^-**1** complex (blue) superimposed with the structure of EV71 3C^pro^-AG7088 complex, revealing large conformational changes of a catalytically important loop 141-147 aa. Fragment **1** (red) and AG7088 (green) are shown with stick models. (B) Zoom in view of the 141-147 aa loop. Residues on the loop are shown with stick models. (C) Molecular surface of 3C^pro^-**1** complex. AG7088 is modeled to substrate binding pockets of 3C^pro^ via structure superimposition. Due to **1** binding induced conformational changes, S1 pocket collapsed and it cannot accommodate the P1 residue. Substrate pockets S1-S5 are highlighted in yellow, the cryptic pocket on leaving group side S’ is highlighted in cyan. (D) Molecular surface of 3C^pro^-AG7088 complex. AG7088 occupies substrate pockets S1-S5, highlighted in yellow; the residues forming the cryptic pocket on leaving group side S’ are highlighted in cyan.

### Conformational rearrangement of the active site induced by covalent fragment binding

To our knowledge, the cryptic S’ pocket identified in our 3C^pro^-fragment complexes has not been observed in the APO 3C^pro^ structures or other 3C^pro^-inhibitor complex structures in the public database. By superimposing our structures with an EV71 3C^pro^-AG7088 complex (PDB ID; 3R0F), we observed a large conformational rearrangement of a catalytically important loop 141-147 aa. This loop harbors the catalytically critical residue C147 and constitutes the upper wall of the S1 pocket. While the 141-147 aa loop remains flexible in the absence of ligand, the binding of substrate (or inhibitor) can hold this loop in the catalytically active conformation, allowing residues G145 and C147 to form the oxyanion hole for the binding of tetrahedral intermediate anion.

The unusual conformation of the 141-147 aa loop in 3C^pro^-**1** and 3C^pro^-**2** structures led to a series of conformational rearrangements: (1) Residue C147 side chain tilted towards the leaving group side of the active site. Comparing to EV71 3C^pro^-AG7088 complex, displacement of the nucleophilic Sγ atom in our structures was 6.4 Å, indicating the geometry of the Ser-His-Asp catalytic triad was disrupted. Displacement of the NH group of G145 was 4.6 Å and displacement of NH of C147 was 1.8 Å, indicating the oxyanion hole could not form. (2) The upper wall of the S1 pocket (the most important pocket for substrate recognition) collapsed, and the size of the pocket became too narrow to accommodate the P1 residue. (3) The leaving group side pockets S1’ and S2’ disappeared, and a previously unobserved cryptic S’ pocket was generated. Residues constituting the cryptic S’ pocket involve I104, T106, H108, M019, M112, V114, F140, T142, A144, G145 and Q146.

### The hit acrylamide fragments inhibit EV71 3C^pro^ and SARS-CoV-2 M^pro^ activity *in vitro*

To investigate which structural features of the thiazole-acrylamide fragments are key to binding, we tested analogues **5**, **6** and **7** for their kinetic binding profiles against 3C^pro^ using qIT (Table 1). Benzothiazole **5** retained potency (REF = 13.5) with a similar kinetic profile to thiazole **1**, further indicating that the thiazole motif drives the binding. Conversely, the *N*-H acrylamides **6** and **7** reacted with 3C^pro^ >50 times more slowly than the parent acrylamide **1**, indicating that alkyl functionalization of the amide nitrogen is required for efficient binding. Indeed, tolerance of the cyclopropane ring of acrylamide **5** indicates that larger substituents may be introduced here and based on the crystal structures of fragments **1** and **2**, this represents a suitable vector for fragment growth towards the canonical binding groove.

**Table 1.**
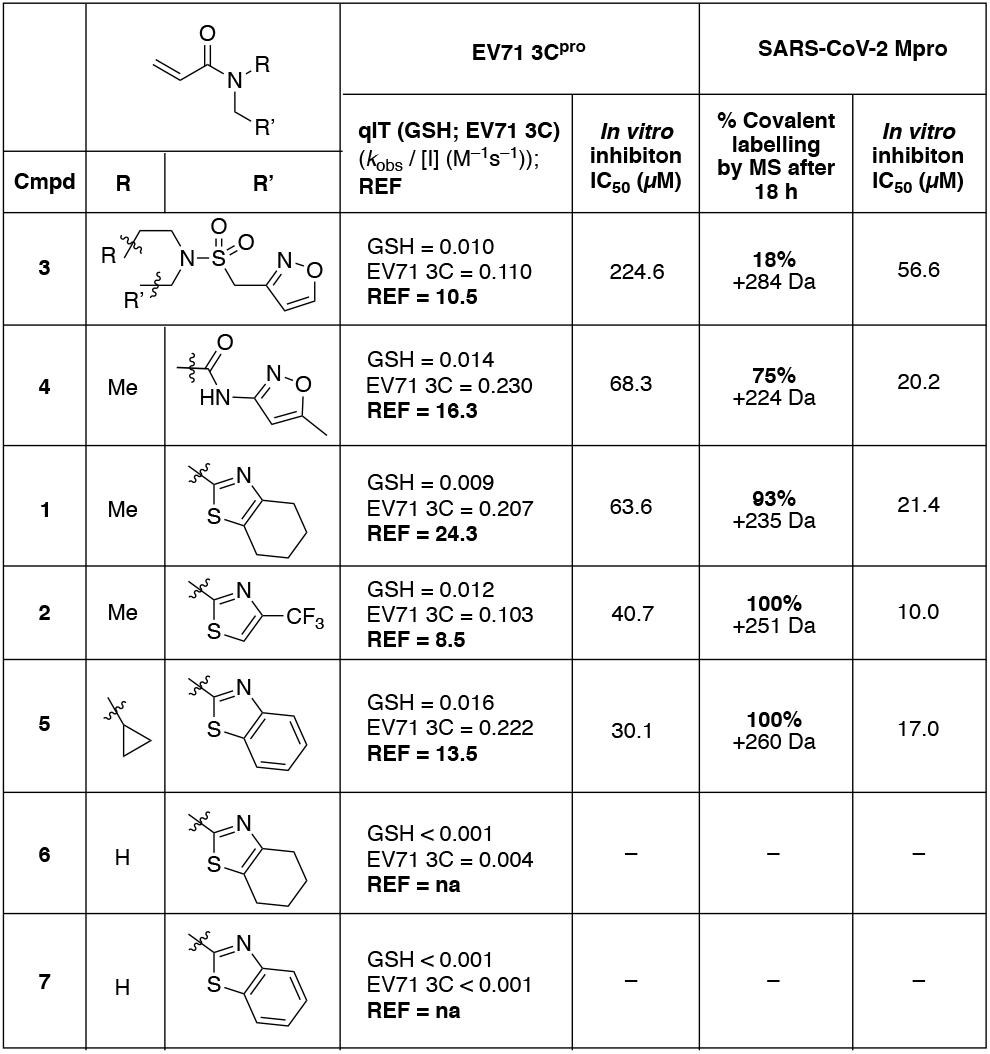
Biochemical characterization of acrylamide fragments.

Next we employed a fluorogenic peptide-based 3C^pro^ protease activity assay to assess biochemical potency of the acrylamides. In accordance with the qIT data, acrylamides **1**-**5** all demonstrated concentration dependent inhibition of 3C^pro^ with modest potency (IC_50_ = 30-230 μM) that is typical of unoptimized fragments.

With the emergence of SARS-CoV-2 as a threat to global public health, we sought to determine if our 3C^pro^-selective fragments could be re-purposed towards this second plus strand RNA virus. Given that 3C^pro^ and M^pro^ are both cysteine proteases that share similar chymotrypsin-folds, we hypothesised that our acrylamide fragments might also be effective against M^pro^. The first examples of irreversible SARS-CoV-2 M^pro^ inhibitors have already emerged,^19,26,27^ but novel acrylamide-based M^pro^ inhibitor scaffolds remain highly desirable. Accordingly, we incubated each fragment with SARS-CoV-2 M^pro^ and used intact protein mass spectrometry to check for covalent modification (Table 1 and Figure S3). Encouragingly, acrylamides **1**, **3** and **4** showed partial modification while fragments **2** and **5** both labelled M^pro^ to completion. Using an M^pro^ activity assay, we validated these results and found that the inhibitory potency against M^pro^ is overall greater than 3C^pro^ (IC_50_ = 10-60 μM), with acrylamides **2** and **5** being the most potent against either proteases. Although other covalent fragment inhibitors of M^pro^ have recently been disclosed,^19^ to our knowledge, our fragments represent the first examples of acrylamide-based fragment inhibitors of M^pro^. Acrylamide-based electrophiles offer low pharmacological risk as indicated by their widespread clinical use, emphasizing the development potential of fragments **2** and **5**.

### Mechanism of the fragment efficacy against M^pro^

To reveal the inhibitory mechanism of these fragments against M^pro^, we determined the crystal structure of M^pro^ with **2** and with **5** (Figure 6). The crystals of M^pro^-**5** complex diffracted to 2.3Å, had a P212121 space group and contained one M^pro^ dimer in the asymmetric unit (ASU). Structural comparison of two monomers in ASU gave an r.m.s.d. of 0.73Å. The crystals of the M^pro^-**2** complex diffracted to 1.8Å, had a C2 space group and contained a single M^pro^ molecule in the ASU. It formed a typical M^pro^ dimer with the symmetry mate (-x, y, -z), suggesting two M^pro^ protomers have identical conformation.

**Figure 6.**
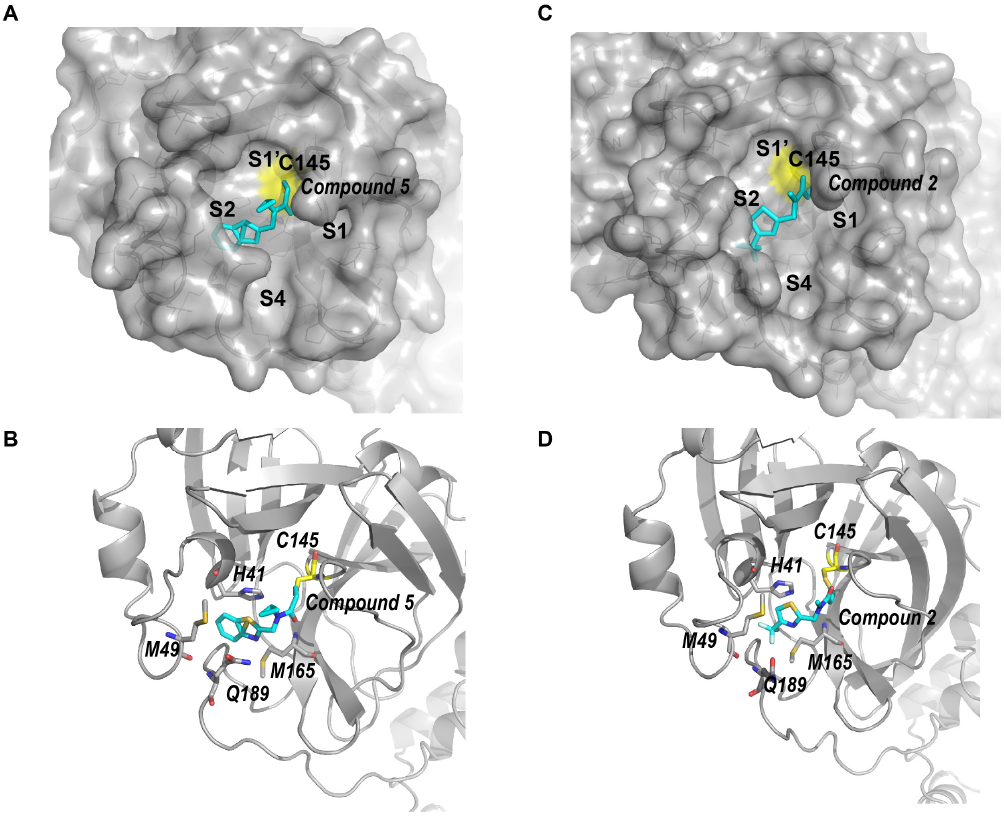
Structure of SARS-CoV-2 M^pro^ complexed with compound **2** & **5**. (A) Surface plot of SARS-CoV-**2** M^pro^ complexed by compound **5** (cyan). Compound 5 occupies pockets S1’ and S2. (B) Ribbon model of SARS-CoV-2 M^pro^-**5**. Residues surrounding the benzothiazole moiety of **5** are shown with stick model. (C) Surface plot of SARS-CoV-2 M^pro^ complexed with **2** (cyan). The trifluoromethyl thiazole moiety of **2** occupies the S2 pocket. (D) Ribbon model of SARS-CoV-2 M^pro^-**2**. Residues surrounding the trifluoromethyl thiazole moiety are shown with stick model.

We identified electron density of compound **5** connecting to the active site cysteine C145 in both M^pro^ monomers in ASU. For the M^pro^-**2** complex, we observed electron density of **2** connecting to the active site residue C145. We generated the polder maps for the above structures with the compound **2** or **5** omitted (Figure S4). Positive densities clearly delineated the structure of compounds **2** and **5**, confirming the presence of the fragments.

In the active site, the acrylamide moiety of **5** forms a covalent bond with C145 (Figure 6 A&B). While the R’ group (benzothiazole) of **5** is accommodated in the deep S2 pocket of M^pro^, the cyclopropane group is exposed to solvent. The benzothiazole/S2 pocket interaction is mainly hydrophobic, involving residues H41, M49, Q189 and M165. The stacking of the H41 imidazole side chain with the benzothiazole moiety stabilizes the fragment.

Similarly, the acrylamide moiety of **2** is covalently linked to residue C145 (Figure 6 C&D). Owing to the high resolution of M^pro^-**2** structure and unambiguous electron density for the fragment, we were able to build the fragment more accurately. We measured the length of the S-C bond between C145 and compound **2** to be 1.8 Å, very close to the average length of single S-C bond, 1.82Å.^28^ The trifluoromethyl thiazole moiety of compound **2** is also accommodated by the S2 pocket. While the trifluoromethyl group touches the apex of the pocket and the thiazole ring π-stacks with the side chain of H41. Both 2 and 5 occupy only the S1’ and S2 subsites, implying substantial opportunity to develop these fragments.

While picornavirus 3C^pro^ functions as a monomer, coronavirus M^pro^ is an obligate dimer. An additional C-terminal domain in M^pro^ stabilizes dimerization and the dimerization interface is essential to maintain the active conformation. We next investigated the oligomerization state of the five M^pro^-fragment complexes using size-exclusion chromatography (Figure S5). As expected, M^pro^-**2**, M^pro^-**3** and M^pro^-**4** eluted as dimers with calculated molecular masses of 45.7kDa, 45.7 kDa and 47.9 kDa, respectively. Interestingly, however, M^pro^-**5** eluted as a monomer. The calculated molecular mass of M^pro^-**5** is 25.3 kDa, whereas the theoretical molecular mass of M^pro^ monomer is 33.7 kDa. The retention volume of M^pro^-**1** lies between the monomeric and dimeric forms, with a calculated molecular mass of 37.6 kDa. This suggests the dimerization was partially impaired.

### Unique inhibitory mechanism of compound 5

To further validate the effects of **5** on M^pro^ dimerization, we tested the oligomerization state of two M^pro^ mutants, C145A and C156W in the absence and presence of ligand. As well as the active site cysteine (C145), C156 is also surface exposed and potentially reactive so the C156W mutant served as a control mutation. Indeed, mutant C156W behaved similarly to the wild-type enzyme: apo-M^pro^ C156W eluted as dimers in size-exclusion chromatography, whilst labelling with **5** retarded M^pro^ elution to that expected for a monomer (Figure 7A). By contrast, mutant C145A remained dimeric irrespective of the presence of **5**. Given mutation C145A prevents the labelling of the active site cysteine, these results clearly indicate that the labelling of C145 by acrylamide **5** is solely responsible for dimer disruption.

**Figure 7.**
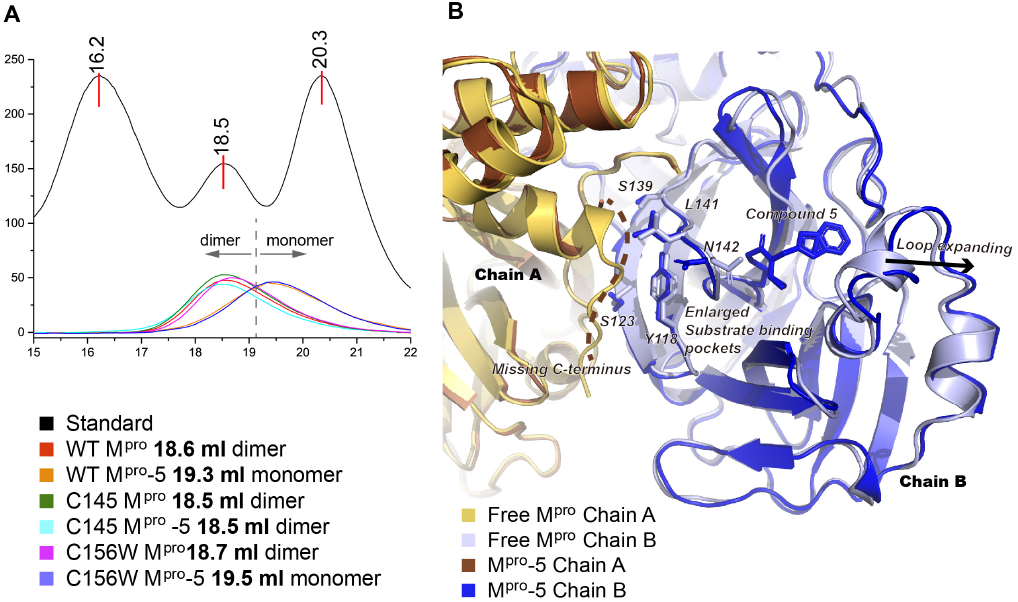
Inhibitory mechanism of **5**. (A) Size-exclusion chromatography analyses two M^pro^ mutants C145A and C156A, the unliganded and labelled with **5**. Elution volume of standards and M^pro^ variants are indicated. (B) Structural comparison of the unliganded SARS-CoV-2 M^pro^ dimer (PDB id: 6Y2E) with M^pro^-**5** dimer. The labelling with **5** expanded the substrate binding pockets. The conformation of several residues (shown with stick model) at the dimer interface were altered. Loops expanding outward are indicated with the arrow. The missing C-terminal region of M^pro^-**5** chain A is indicated with the dashed line.

Our crystallographic data provides further insights into the inhibition mechanism. Although M^pro^-**5** forms dimers in crystal lattices, these are notably different from authentic M^pro^ dimers: (1) Most published M^pro^ dimers have 2-fold symmetry between two protomers, but M^pro^-**5** protomers exhibit marked difference, r.m.s.d =0.73Å. The structure of each M^pro^-**5** protomer is also notably different from the free enzyme (PDB id: 6Y2E), r.m.s.d= 0.71-0.78 Å. In this regard, M^pro^-**2** is more similar to the free enzyme. The structure of M^pro^-**2** dimer has 2-fold symmetry and each protomer is highly similar to the free enzyme, r.m.s.d =0.22 Å. (2) The binding of **5** enlarged the substrate binding pockets and affected the nearby regions (Figure 7B). Comparing to the free enzyme, the loops surrounding **5** expanded to make room for benzothiazole motif. This induced conformational changes of several residues and regions at the dimerization interface. In the chain A of M^pro^-5 dimer, the extreme C-terminal region at the dimer interface went disordered, which was likely caused by the labelling of compound **5** on C145. The fragment induced conformational alterations may contribute to the destabilization of M^pro^-**5** dimers. In summary, we found that **5** has at least two mechanisms of action to inhibit M^pro^: (1) covalently linking to the catalytically important cysteine and occupying the substrate binding pockets; (2) destabilizing the dimerization of M^pro^.

In conclusion, we have identified acrylamide fragments that target both the EV71 3C^pro^ and SARS-CoV-2 M^pro^, and inhibit their activity by covalently reacting with their catalytic cysteines. Importantly, some of these hit fragments cause profound structural rearrangements of each protease: in the case of EV71 3C^pro^ a new subsite pocket is formed at the expense of the normal active site architecture whilst with M^pro^ key structural features required for dimerization are distorted preventing formation of the active dimeric unit. The discovery of these conformational change-based mechanisms of action on covalent fragment binding demonstrates the utility of solution-based screening methodologies as an alternative to crystallographic fragment screening in which structural rearrangements are unlikely. These hit ligands also provide excellent candidates for development of potent protease inhibitors.

## ASSOCIATED CONTENT

### Supporting Information

The Supporting Information is available free of charge on the ACS Publications website.

Experimental details, supporting Figures S1-5 and Table S1 (PDF)

## Author Contributions

The manuscript was written through contributions of all authors.

