## Supplementary Information for "Acrylamide Fragment Inhibitors that Induce Unprecedented Conformational Distortions in Enterovirus 71 3C and SARS-CoV-2 Main Protease"

- **J.** .......... Error! Bookmark not defined.
- **K.** .......... Error! Bookmark not defined.
- **L.** .......... Error! Bookmark not defined.
- **M.** .......... Error! Bookmark not defined.
- **N.** .......... Error! Bookmark not defined.
- **O.** .......... Error! Bookmark not defined.
- **P.** .......... Error! Bookmark not defined.
- **Q** .......... Error! Bookmark not defined.
- **R.** .......... Error! Bookmark not defined.

**Supplementary Reference** .......... 11

### Supplementary Results

#### Supplementary Figures

Figure S1: Rate enhancement factor analysis
Figure S2: qIT and intact protein mass spectrometry of validated ligands
Figure S3: Intact protein mass spectroscopy of hit ligands
Figure S4: The polder maps of fragments linked to Mpro
Figure S5: Size-exclusion chromatographic analysis of various Mpro-fragment complexes
Figure S5:
Figure S6:

#### Supplementary Tables

Table S1: Data collection and refinement statistics

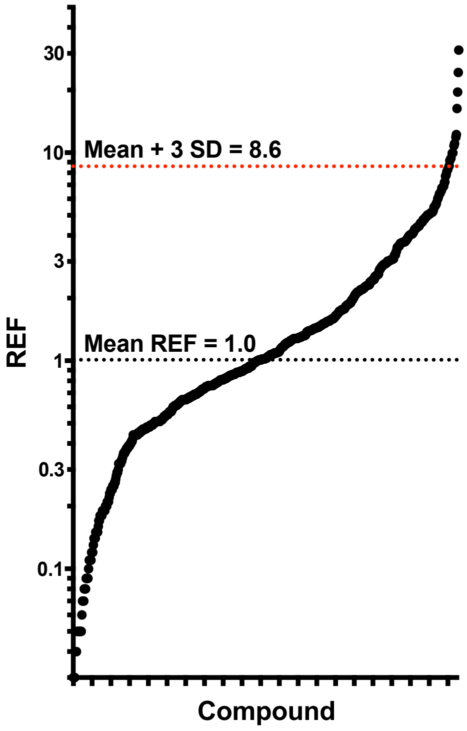

**Supplementary Figure S1 | Rate enhancement factor analysis.** Compounds (0.5 mM) were reacted with EV71 3C or GSH (5 µM) at pH 8.0 and the rate of reaction ($v$) determined by qIT. The rate enhancement factor (REF) was calculated: REF = $v$(3C)/$v$(GSH). The 639 fragments for which a REF could be calculated are plotted, highlighting the geometric mean REF (1.0) and the standard deviation (SD = 2.5). Hits were defined as REF > geomean + 3SD.

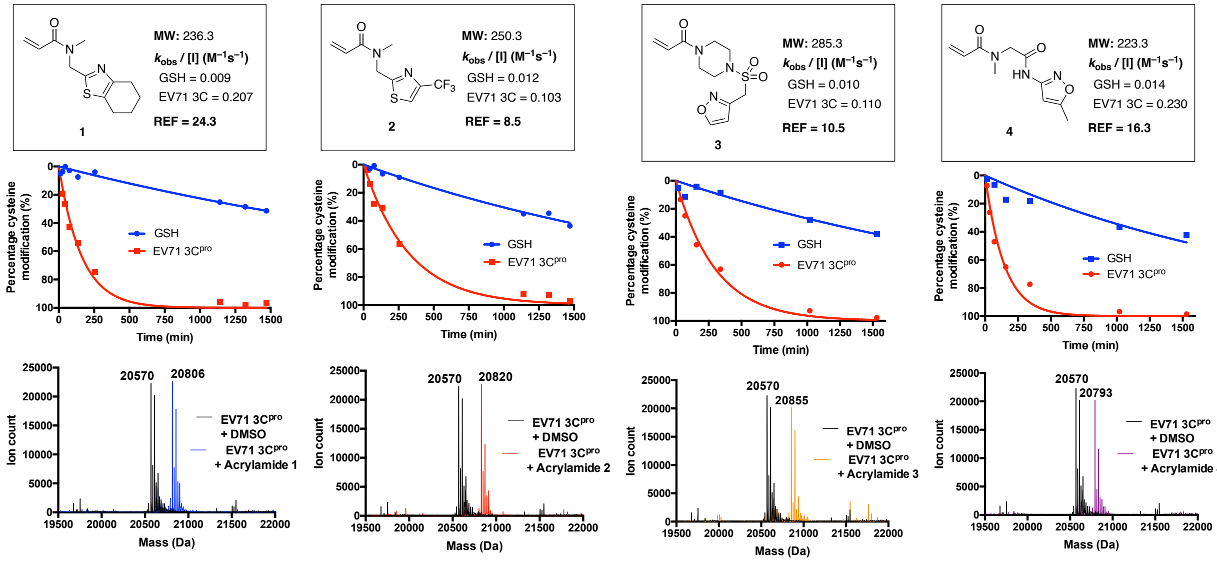

**Supplementary Figure S2 | qIT and intact protein mass spectrometry of hit acrylamides against EV71 3c.** For each hit fragment the qIT profile and intact protein mass spectrum are shown. For qIT data: Acrylamides 1-4 (0.5 mM) were reacted with EV71 3C or glutathione (5 µM) in pH 8.0 HEPES buffer and the reaction progress was followed by discrete measurements of thiol concentration using the fluorogenic probe CPM . Fluorescence intensity is converted into percentage cysteine modification by normalizing to DMSO control = 0%, no thiol = 100%. Rate constants are derived from exponential regression analysis. For intact protein mass spectrometry: EV71 3C (5 µM) was treated with

DMSO or acrylamides 1-4 (0.5 mM) and incubated at rt for 750 minutes. Intact protein mass spectra of each acrylamide reaction is overlaid with DMSO control, showing mono-modification in each case.

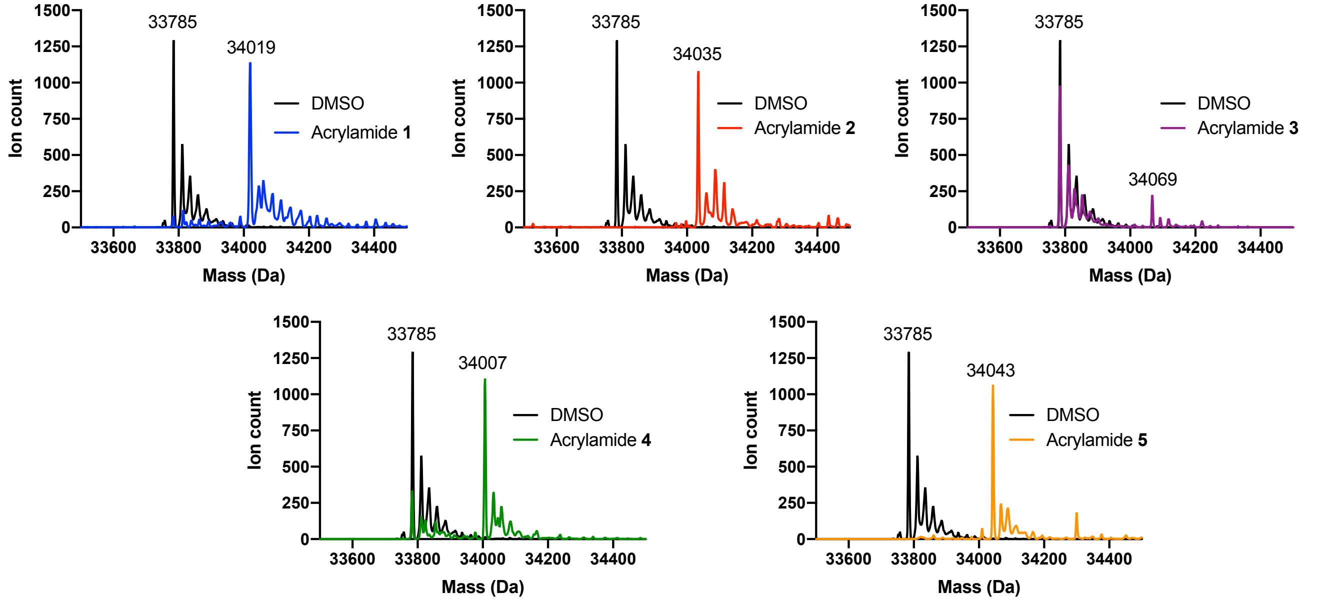

**Supplementary Figure S3 | Intact protein mass spectrometry against SARS-CoV-2 Mpro**. SARS-CoV-2 Mpro (5 μM) was treated with DMSO or acrylamides 1-5 (0.5 mM) and incubated at rt for 1000 minutes. Intact protein mass spectra of each acrylamide reaction is overlaid with DMSO control.

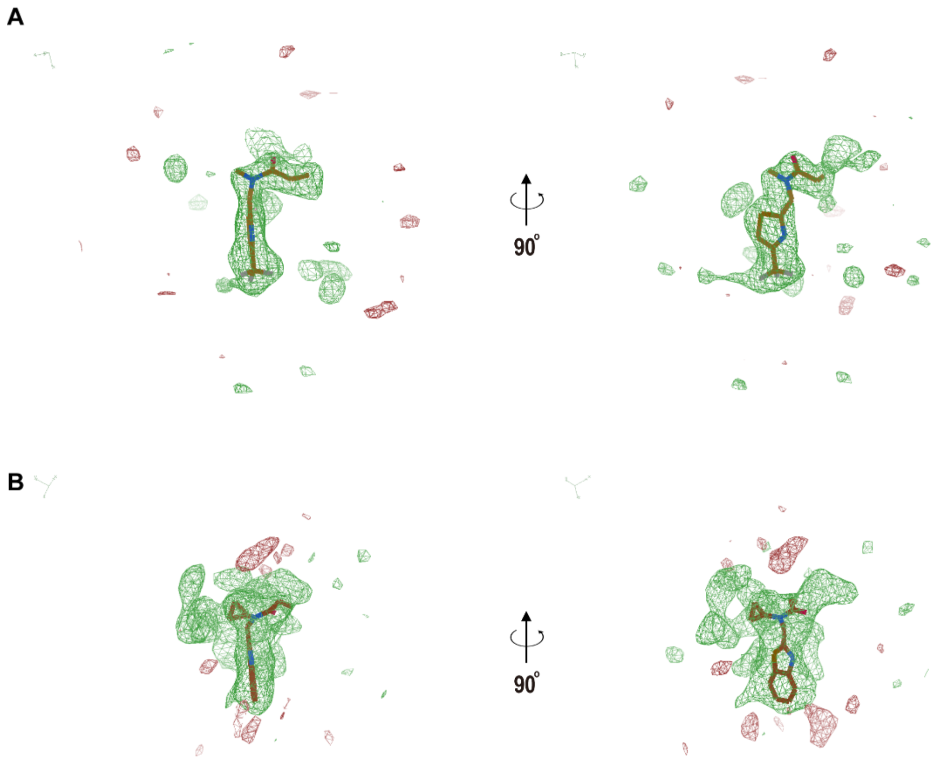

**Supplementary Figure S4 | The polder maps of fragments linked to Mpro**. The polder maps of the crystal structure of SARS-CoV-2 Mpro complexed by compound 2 (A) and 5 (B). The map was calculated using the software phenix.polder. The omitted region was the fragment itself. The positive and negative difference (mFobs-DFmodel OMIT) is shown in green and red densities with the contour level± 1.5 σ.

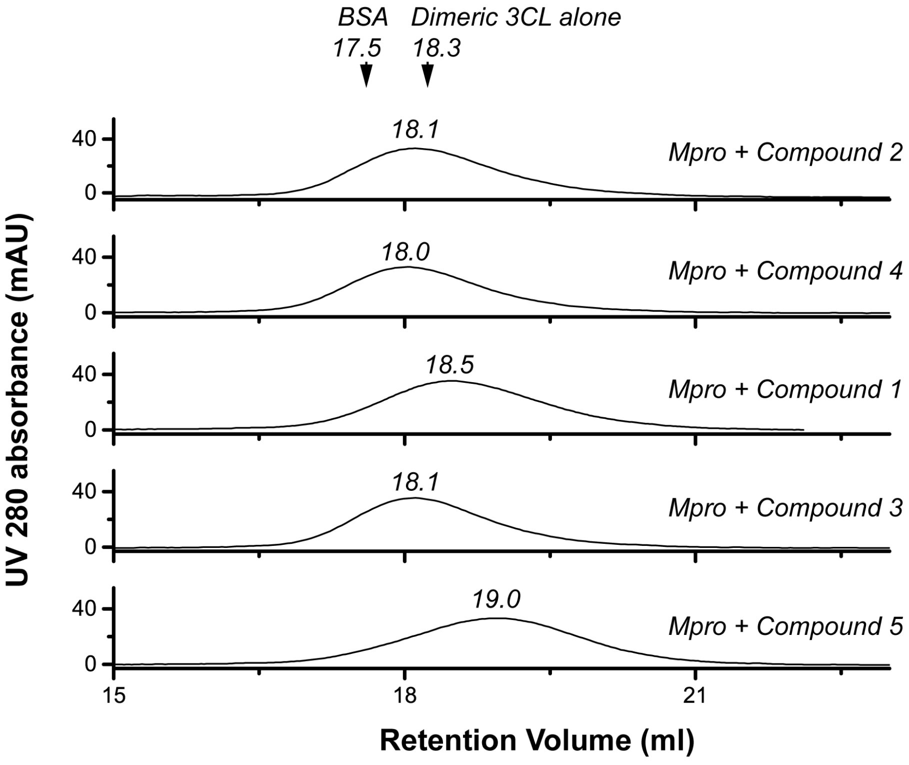

**Supplementary Figure S5 | Size-exclusion chromatographic analysis of various Mpro-fragment complexes.** Mpro was pre-incubated with compound 1-5 respectively and the complexes were loaded to Superdex 200 10/300GL column. The retention volumes of the Mpro-fragment, BSA and the unliganded dimeric Mpro are indicated.

**Tables S1 | Data collection and refinement statistics**

| | **EV71 3C$^{pro}$ compound 2 (PDB ID:7CNI)** | **EV71 3C$^{pro}$ compound 1 (PDB ID:7CNH)** | **SARS CoV-2 M$^{pro}$ compound 2 (PDB ID:7CNK)** | **SARS CoV-2 M$^{pro}$ compound 5 (PDB ID:7CNJ )** |
|---|---|---|---|---|
| **Data collection** | | | | |
| **Space group** | $P2_12_12_1$ | $P2_12_12_1$ | C2 | $P2_12_12_1$ |
| **Cell dimensions** | | | | |
| **a, b, c (Å)** | 59.77, 70.50, 86.00 | 59.78, 70.76, 85.46 | 114.40, 53.70, 44.80 | 67.90, 101.40, 103.30 |
| **α, β, γ (°)** | 90.00, 90.00, 90.00 | 90.00, 90.00, 90.00 | 90.00, 101.30, 90.00 | 90.00, 90.00, 90.00 |
| **Resolution (Å)** | 49.08 (1.23) | 23.57 (1.41) | 48.47 (1.81) | 49.45 (2.30) |
| $R_{sym}$ | 0.04 (0.63) | 0.06 （0.86） | 0.05 (0.79) | 0.05(0.84) |
| $I / \sigma I$ | 38.50 (1.34) | 42.85(2.44) | 25.44(0.89) | 27.17(0.90) |
| **Completeness (%)** | 98.9 (39.50) | 99.20 (99.80) | 97.30 (37.10) | 99.90 (99.70) |
| **Redundancy** | 3.31 (0.74) | 13.72 (12.94) | 4.41(0.94) | 6.60(6.38) |
| **Refinement** | | | | |
| **Resolution (Å)** | 49.08 (1.23) | 23.12 (1.40) | 23.49 (1.80) | 45.96 (2.30) |
| **No. reflections** | 168929 | 71281 | 20040 | 32303 |
| $R_{work}$ / $R_{free}$ | 0.2333/0.2566 | 0.1838/0.197 | 0.2014/0.2425 | 0.2081/0.2698 |
| **No. atoms** | | | | |
| **Protein** | 5638 | 5685 | 4668 | 4687 |
| **Ligand/ion** | 27 | 33 | 24 | 54 |
| **Water** | 393 | 512 | 111 | 81 |
| ***B*-factors** | | | | |
| **Protein** | 18.73 | 23.57 | 51.68 | 55.55 |
| **Ligand/ion** | 18.22 | 22.36 | 87.94 | 69.37 |
| **Water** | 25.99 | 33.52 | 47.22 | 52.48 |
| **R.m.s. deviations** | | | | |
| **Bond lengths (Å)** | 0.012 | 0.007 | 0.005 | 0.007 |
| **Bond angles (°)** | 1.22 | 0.99 | 0.77 | 0.97 |
| **% residues in favored regions, allowed regions, outliers in Ramachandran plot** | 97, 3, 0 | 97, 3, 0 | 98, 2, 0 | 96, 4, 0 |

*Values in parentheses are for highest-resolution shell.

### Supplementary Methods

#### A. Library design

Acrylamide fragments were synthesized from amine precursors (see *Efficient and Facile Synthesis of Acrylamide Libraries for Protein-Guided Tethering* for full synthetic details)[1] or purchased directly from commercial supplier and used as supplied. The physiochemical properties of the final library were calculated using DataWarrior [2] and found to broadly comply with the 'Rule of 3' guidelines [3], with the recommended relaxed restriction on number of H-bond acceptors[4] (Supplementary Fig. S2b). Compounds **1**–**7** were purchased from Enamine.

#### B. Plasmid construction

The DNA encoding the EV71 3C proteinase (1–183 aa) was amplified and inserted into pET28a (Novagen). The resulting plasmid encodes an N-terminal His-tagged EV71 3C proteinase. Amino acid sequence of the N-terminal extension is MGSSHHHHHHSSGLVPRGSHM, which contains the 6×His and a thrombin cleavage site. Plasmid expressing a EV71 3C mutant harboring mutation H133G was generated using site-directed mutagenesis (QuikChange™). The sequence of the plasmid was verified by DNA sequencing.
The plasmid for expressing the dimeric Mpro was constructed according to a previous study [7]. Briefly, the gene of SARS-CoV-2 Mpro (1-306aa, NCBI Reference Sequence: YP_009742612.1) was synthesized (Sangon Shanghai, China) and inserted into the PGEX-6p-1 plasmid (GE Healthcare). The N-terminal portion of the construct contains the auto cleavage site of Mpro SAVL**Q/S**GFRK derived from the boundary of nsp4/nsp5 of the polyprotein precursor ORF1ab. The C-terminal portion contains a PreScission cleavage site VTF**P/G**P and a 6xHis tag. This design is important for selecting only the folded and active Mpro that could auto cleave its own N-terminal site. Mutations were introduced to the plasmid expressing the dimeric Mpro using the QuickChange methods to express mutants C145A and C156W.

#### C. EV71 3C expression and purification

Expression and purification of recombinant 3C proteinase followed the protocol established previously.[5],[6] Briefly, plasmid was transformed into Rosetta™ (DE3) Competent Cells (Novagen). Bacteria culture was grown in lysogeny broth medium (LB) at 37°C to a density $OD_{600}$ ~ 1.0. Isopropyl-D-1-thiogalactopyranoside (IPTG) was then added to culture (final concentration of 0.5 mM) to induce expression, and the shaking was continued for another 20 h at 18°C. Bacterial cells were collected by centrifugation at 2,991g, and resuspended in the lysis buffer containing 50 mM Tris-HCl (pH 8.0), 150 mM NaCl and 20mM imidazole. The bacterial cells were disrupted by ultrasonication. Cell debris was removed by centrifugation at 20,000 g for 1h. The supernatant was loaded to Ni-nitrilotriacetic acid resin (GE Healthcare, USA) pre-equilibrated with the lysis buffer. Nonspecific contaminants were removed by washing the resin with 10x bed volume of the wash buffer containing 50 mM Tris-HCl (pH 8.0), 150 mM NaCl, and 20 mM imidazole. Recombinant 3C proteins was finally eluted with the elution buffer containing 50 mM Tris-HCl (pH 8.0), 150 mM NaCl, and 300 mM imidazole. Thrombin (Sigma) was then used to cleave 6xHis-tag. 3C proteinase was finally purified by gel-filtration (Superdex 200 column, GE Healthcare, USA) pre-equilibrated with the gel-filtration buffer containing 25 mM HEPES (pH 8.0), 150 mM NaCl. Fractions containing 3C proteinase were pooled and concentrated to 10 mg/ml.

#### D. SARS-CoV-2 Mpro and its variants expression and purification

The expression of all SARS-CoV-2 Mpro variants follows the same protocol. Plasmids were transformed to BL21 (DE3) competent cells. The bacterial culture was grown to $OD_{600}$ 0.6-0.8 at 37 ℃ before inducing with 0.5mM IPTG. The shaking of the bacteria continued at 18 ℃ overnight. The bacterial cells were collected by centrifuging at 5,000 rpm; cell pellet was cooled on ice and resuspended in 80 ml lysis buffer (50 mM Tris, 150 mM NaCl, pH 8.0, 0.16% mercaptoethanol, 1% PMSF). The cells were disrupted by ultrasonication on ice and clarified by centrifugation at 20,000 rpm.

(1) To prepare dimeric SARS-CoV-2 Mpro or its mutant, the supernatant was first loaded to Ni-NTA resin and the His-tagged Mpro was eluted with 10x column volume elution buffer (50 mM Tris, 150 mM NaCl, pH 8.0, 300 mM imidazole, 0.16% mercaptoethanol). At this stage, most Mpro lost its N-terminal GST tag via auto cleavage. The His-tagged Mpro was digested with 200 μ g PreScission protease through dialysis overnight at 4 ℃ in the dialysis buffer containing 50 mM Tris, 150 mM NaCl, pH 8.0, 1mM DTT. The Mpro with authentic N- and C-terminus was finally obtained by passing the sample through the GST resin and Ni-NTA resin. Residual Mpro with the uncleaved GST tag, the GST-tagged PreScission protease and the his-tag containing species were all trapped with the resins. The final step of the purification was the Superdex 200 10/300 column equilibrated by the gelfiltration buffer (25 mM

Hepes, 150 mM NaCl, pH 8.0). The peak corresponding to dimeric Mpro was collected. The Mpro was concentrated to ~10 mg/ml before use or stored at -80℃.

(2) The preparation of Mpro mutant C145A follows the similar protocol for preparing dimeric Mpro with one additional step after Ni-NTA purification. Given this mutant is catalytic inactive and cannot auto cleave its own N-terminal tag, wild-type Mpro was added to Ni-NTA elution and the sample was dialyzed overnight to allow proteolytic processing. The subsequent purification steps remained the same. C145A mutant harbors authentic N- and C-terminus and eluted as dimers.

#### E. Rate determination by qIT

*Reaction-plate setup*: Each well of a 384-well (Corning – black, NBS) plate was charged sequentially with 20 µL of TCEP-agarose beads (3x stock (1.5% slurry) and 20 µL of target thiol (3x stock (15 µM) of GSH or EN71 3C(WT)) both in 25 mM HEPES (pH 8.0), 150 mM NaCl buffer and incubated at r.t. for at 1-2 hours to ensure complete reduction. Reactions are started by addition 20 µL of electrophile solution (3x stock, 1.5 mM, 3% DMSO) in 25 mM HEPES (pH 8.0), 150 mM NaCl buffer (final volume = 60 µL, final concentration: thiol = 5 µM, electrophile = 500 µM, 0.5% w/v TCEP-agarose beads). Timer is started. After mixing, TCEP-agarose was pelleted by centrifugation of the plate (1,000 rpm, 5 min).
*CPM Quench***:** At a series of time points (typically 7, 15, 30, 60, 120, 240, 1100, 1300, 1440 minutes) 3 µL aliquots of each reaction were quenched into a 384-well fluorescence plate (Corning – black, NBS) where each well was charged with 27 µL of CPM solution (1.39 µM) in quench buffer (25 mM HEPES (pH 7.5), 150 mM NaCl buffer) (final concentration: thiol = 0.5 µM, CPM = 1.25 µM). Quenching was typically performed in singlet and quench plates are incubated at room temperature for 60 minutes before fluorescence intensity (excitation/emission: 384/470 nm) were measured on a Clariostar plate reader.
*Analysis*: All analysis was performed using Prism software (GraphPad). For each time point, the average fluorescence of the [DMSO – thiol] is subtracted from each sample. Then each sample is divided by the average of the [DMSO + thiol] controls (which have also had the [DMSO – thiol] subtracted). This normalized fluorescence (NF) value is converted to percentage modified cysteine for each reaction (percentage modified cysteine = (1-NF)/-100) and is plotted against time. A one phase exponential decay was fitted to each plot, yielding a pseudo first-order rate constant.

#### F. Intact-protein mass spectrometry

EV71 3C or SARS-CoV-2 Mpro (5 µM) were reacted with acrylamide ligands at 500 µM or DMSO in 25 mM HEPES, 150 mM NaCl buffer (pH 8) containing 1% DMSO for 750-1000 minutes. Excess ligand was subsequently removed by serial dilution and concentration (×5) into the ammonium bicarbonate (30 mM) using Amicon Ultra-0.5 centrifugal filter devices (MW cut-off = 10 kDa).
Protein-mass data were obtained on a Waters micromass LC-TOF mass spectrometer in positive ion mode using electrospray ionization. Samples were chromatographed using an Acquity UPLC protein BEH C4 column (2.1 mm × 50 mm, 2.7 µm) with a flow rate of 30 µL/min. The injection volume was 5 µL of protein solution (5 µM) in 30 mM ammonium bicarbonate buffer containing 0.5% formic acid and 20% MeOH. The gradient used was 100% mobile phase A (0.1% formic acid in water/MeOH (19:1)) ramping linearly to 90% mobile phase B (0.1% formic acid in MeOH) over 5 minutes. The spectra were acquired from 500 to 2500 Da and deconvolution was performed with MassLynx (Waters), using the maximum entropy algorithm, over a mass range of 800 to 1200 Da with a mass step of 0.75.

#### G. *In vitro* protease inhibition assay

The half maximal inhibitory concentration (IC50) of each cysteine-reactive fragment was measured using an *in vitro* proteinase assay described previously[3].

(1) For EV71 3C inhibition assessment, each reaction mixture (100µL) contained 25mM HEPES (pH 7.0), 150 mM NaCl, 8µM WT 3C proteinase and the increasing amount of cysteine-reactive fragment or AG7088 as the positive control. The mixtures were pre-incubated at 25°C for 12 h in 96-well plate (Greiner), allowing the labeling reaction to complete. Then, 40 µM fluorescent peptide substrate (Dabcy-N'-lRTATV**Q↓G**PSLDFE-C'-Edans, harboring the a 3C cleavage site), harboring the junction sequence between 3B/3C of EV71 polyprotein precursor, was added to the reaction mixtures. Relative fluorescence unit (RFU) was recorded at 490 nm (emission) when the excitation wavelength was set to 340nm. The RFU values was then converted to substrate concentration using the Edans standard calibration curve that defined the relationship between RFU and concentration. The initial velocities of 3C catalyzed reaction was calculated by averaging at least three measurements.

(2) For SARS-CoV-2 Mpro inhibition assessment, each reaction mixture (100μL) contained 25mM HEPES (pH 7.0), 150 mM NaCl, 1μM WT 3C proteinase and the increasing amount of cysteine-reactive fragment (up to 200μM). The mixtures were pre-incubated at 25°C for 12 h in 96-well plate (Greiner), allowing the labeling reaction to complete. 40 μM fluorescent peptide substrates (Dabcyl-N'-KTS AVL**Q↓S**GFRKME-C'-Edans) harboring the Mpro cleavage site between nsp4/nsp5 juncture of SARS-CoV-2 polyprotein precursor ORF1a, was added to the reaction mixtures. Relative fluorescence unit (RFU) was recorded at 490 nm (emission) when the excitation wavelength was set to 340nm. The RFU values was then converted to sub-strate concentration using the Edans standard calibration curve that defined the relationship between RFU and concentration. The initial velocities of the Mpro catalyzed reaction was calculated by averaging at least three measurements.

To calculate IC50 for either EV71 3C or SARS-CoV-2 Mpro inhibition by those inhibitors, reaction velocity was plotted as the function of the inhibitor concentration. The data was analyzed using GraphPad software with the equation [inhibitor] vs. response (three parameters), Y=Bottom + (Top-Bottom)/(1+(X/IC50)).

#### H. Crystallization and structure determination for EV71 3C

EV71 3C mutant H133G was used for crystallographic study. The protein was concentrated to 1 mg/ml in a buffer containing 25 mM HEPES (pH 8.0) and 150 mM NaCl. The fragment power was dissolved in 100% dimethyl sulfoxide (DMSO) with the concentration of 50 mM. To prepare 3C-fragment complex, protein and fragment were mixed with a molar ratio of 1:22. The mixture was incubated at 25 °C for 24 h to allow labeling. Minor precipitation was visible after the incubation; the mixture was then clarified by centrifugation at 15,294 g. To remove unbound fragment, the clarified mixture was first concentrated with 15 ml centrifugal filter (10 kDa cutoff) and then 10x volume fresh buffer was added. The procedure was repeated 3 times to ensure the complete removal of the unbound. Finally, 3C-fragment complex was concentrated to ~10 mg/ml before crystallization trials. To grow crystals, 1μl 3C-fragment sample was mixed with 1μl reservoir buffer, and the drops were incubated at 18 °C in a vapor diffusion system. The optimized crystallization buffer for 3C-337 complex contains 6% Tacsimate pH 7.0, 14% PEG3350; the optimized crystallization buffer for 3C-338 complex contains 4% Tacsimate pH 7.0, 12% PEG3350.
X ray diffraction data of 3C-**1** and 3C-**2** crystals were collected at 19U beamline of Shanghai Synchrotron Radiation Facility and PXIII beamline of Swiss Light Source, Paul-Scherrer Institute, Switzerland. Diffraction data was processed using XDS Package. [7] 3C-**1** and 3C-**2** crystals diffracted the X ray to 1.23 Å and 1.40 Å. They shared the space group of P212121and contained two 3C copies per asymmetric unit. Both structures were solved by molecular replacement using EV71 3C H133G (3R0F) as the searching model using the software Phaser for MR.[8] Loop 141-147 aa were manually built using the software Coot. The structures were refined using the software Phenix Refine.[9] Parameters of data collection and structure refinement are summarized in Table S1.

#### I. Crystallization and structure determination of SARS-CoV-2 Mpro complexed by inhibitors

To crystallize SARS-CoV-2 Mpro complexed by compound 2 and 5. The Mpro was concentrated to 1 mg/ml in a buffer containing 25 mM HEPES (pH 8.0) and 150 mM NaCl. The fragment was dissolved in 100% DMSO with the concentration of 50mM. The Mpro and the fragment were mixed with a molar ratio of 1:22 and was incubated at 25 °C for 18 hrs to allow labeling. The mixture was then clarified by centrifugation at 15,294 g. To remove the unbound fragment, the mixture was concentrated with a 15 ml centrifugal filter (Millipore, 10kDa cutoff) and then diluted with 10x volume fresh buffer. The procedure was repeated 3 times to ensure complete removal of the unbound inhibitors. Finally, Mpro-fragment complex was concentrated before crystallization screening. Mpro-**2** was concentrated to ~ 4.5mg/ml and Mpro-**5** was concentrated to ~ 2.1mg/ml.
To grow crystals, 1μl Mpro-fragment complex was mixed with 1μl reservoir buffer, and the drops were incubated at 18 °C in a vapor diffusion system. The optimized crystallization buffer for Mpro-**2** complex is 0.2 M Ammonium sulfate, 23% PEG 3350 and 0.1 M Bis-Tris pH 6.9; the optimized crystallization buffer for Mpro-**5** complex is 0.1M Hepes pH 7.5 ，6% 2-Propanol and 16% PEG4,000.
X ray diffraction data of Mpro-2 and Mpro-5 crystals were collected at the protein crystallography beamlines at Shanghai Synchrotron Radiation Facility and at Diamond Light Source. Diffraction data was processed using XDS Package. Mpro-**2** and Mpro-**5** crystals diffracted the X ray to 1.81 Å and 2.30 Å. They have the space group of C2 and P212121. All structures were determined by molecular replacement using the software Phaser for MR. The searching model was a Mpro-inhibitor structure 6LZE. The structures were refined using the software Phenix Refine. Manual model building was conducted using software Coot. Parameters of data collection and structure refinement are summarized in Table S1.

[5] Cui, S. *et al.* Crystal structure of human enterovirus 71 3C protease. *Journal of molecular biology* **408**, 449-461

[6] Wang, J. *et al.* Crystal structures of enterovirus 71 3C protease complexed with rupintrivir reveal the roles of catalytically important residues. *Journal of virology* **85**, 10021-10030

[7] Kabsch, W. XDS. *Acta Crystallogr. Sect. D Biol. Crystallogr.* **2010**, *66* (2), 125–132.

[8] McCoy, A. J.; Grosse-Kunstleve, R. W.; Adams, P. D.; Winn, M. D.; Storoni, L. C.; Read, R. J. Phaser Crystallographic Software. *J. Appl. Crystallogr.* **2007**, *40* (4), 658–674.

[9] Liebschner, D.; Afonine, P. V.; Baker, M. L.; Bunkóczi, G.; Chen, V. B.; Croll, T. I.; Hintze, B.; Hung, L.-W.; Jain, S.; McCoy, A. J.; Moriarty, N. W.; Oeffner, R. D.; Poon, B. K.; Prisant, M. G.; Read, R. J.; Richardson, J. S.; Richardson, D. C.; Sammito, M. D.; Sobolev, O. V.; Stockwell, D. H.; Terwilliger, T. C.; Urzhumtsev, A. G.; Videau, L. L.; Williams, C. J.; Adams, P. D. Macromolecular Structure Determination Using X-Rays, Neutrons and Electrons: Recent Developments in Phenix. *Acta Crystallogr. Sect. D Struct. Biol.* **2019**, *75* (10), 861–877.
